# Metabolic characteristics of taste differences under the soil and hydroponic cultures of sweet potato leaves by using non-targeted metabolomics

**DOI:** 10.1101/2021.02.24.432602

**Authors:** Zhaomiao Lin, Guoliang Li, Hong Zhang, Rongchang Ji, Yongqing Xu, Guochun Xu, Huawei Li, Zhonghua Liu, Wenbin Luo, Yongxiang Qiu, Sixin Qiu, Hao Tang

## Abstract

Sweet potato leaves are consumed as green leafy vegetables in most of the world due to their nutritional and functional values, and the taste characteristics determine their commodity value and consumer acceptance. However, the metabolic composition and formation mechanism of taste quality in its leaves are not clear. In this study, we found that sweet potato leaves under different growing patterns, soil culture and hydroponic culture, which result in different taste quality. In particular, the taste quality in leafy sweet potato was effectively improved under hydroponic culture. Meanwhile, we further profiled metabolites in leaves of sweet potatoes under different growing patterns by using GC–QToF–MS. A total of 200 metabolites were identified, covering most of the metabolic pathways in plants. A comparison of the good taste and poor taste of sweet potato leaves resulted in 71 metabolites related to taste quality formation. In addition, the leaves with poor taste had lower levels of metabolites regarding amino acids metabolism, whereas was accompanied by high levels of metabolites in carbohydrates and secondary metabolism. This study provides new insights into the improvement of taste quality in leafy sweet potato.

## INTRODUCTION

Sweet potato (Ipomoea batatas L.) is an important food crop in the world, with its roots be used for food or raw materials in industrial production. Meanwhile, sweet potato leaves, the young parts of the shoot in sweet potatoes, have been used as leafy vegetables in most tropical and subtropical areas (Islam, 2006). This vegetable-use of sweet potato was called leafy sweet potato. Compared with ordinary leafy vegetables, leafy sweet potato have many advantages. Such as, their yield is high due to can be harvested several times a year (Sun et al., 2016); and their nutritional and functional values were much higher than its root or other leafy vegetables (Dinu et al., 2018; Li et al., 2019; Sun et al., 2014); as well as they have a strong resistance to drought, waterlogging, and barren, etc (Sui et al., 2019).

Taste quality is one of the important quality traits of vegetables. In leafy sweet potato, the taste characteristics determine its commodity value and consumer acceptance (Lee et al., 2007; Ona and Chujoy, 1991). It has been reported that hydroponic culture increased vitamin C, flavone, nitrate contents, and yield of sweet potato leaves compared with soil culture (Chen et al., 2013). In the previous study, compared to soil grown strawberries, the hydroponically grown strawberries were wined preferred by 70% of participants in the sensory analysis (Treftz et al., 2015). We also found that hydroponic culture also can effectively improve the taste quality in leafy sweet potato, especially the cultivars with poor taste under soil culture in the field. However, the metabolic composition and formation mechanism of taste quality in its leaves are not clear.

Metabolomics is the modern instrumental analysis method with high-throughput, high-resolution, and high-sensitivity, to comprehensively study the various features of endogenous metabolites in living organisms (Saito and Matsuda, 2010). For taste quality analysis, metabolomic approaches have yielded new insights into the taste change in the breeding history of tomato (Zhu et al., 2018), the sensory quality assessment of garlic (Liu et al., 2019), exploring the relationships between these substances and tastes of white tea (Yang et al., 2018), and Identification of key taste components in loquat (2020). Therefore, in this study, we used the untargeted metabolomic analysis of sweet potato leaves under different growing patterns, soil culture and hydroponic culture, for exploring the metabolic foundation of different taste quality in sweet potato leaves.

## RESULTS

### Differences of taste quality in sweet potato leaves under different growing patterns

Leafy sweet potato is a special type of sweet potato for vegetables, and the taste quality is the one of major concerned trait for the market and consumer. Leafy sweet potato cultivation is mainly based on traditional land cultivation methods. However, use of hydroponic culture to produce leafy sweet potato has many advantages for shoot growth and its yield increase, because it goes against root formation in nutrient solution (Chen et al., 2013). Hydroponic culture is also beneficial for the pollution-free, standardized, and mechanized production of leafy sweet potato. In our pre-experiment, we compared the taste characteristics of the leaves of 8 different sweet potato varieties under hydroponic culture (H) and soil culture (S) conditions, and found that hydroponic culture can effectively improve the taste quality of sweet potato leaves, especially the cultivars with poor taste under soil culture (Table S1). Thus, hydroponics can be used as a reliable cultivation mode to improve the taste quality of leafy sweet potatoes. In addition, the taste quality of Baisheng under soil culture was poor, while the taste quality of Baisheng under hydroponic culture (Baisheng-S) was significantly improved; and taste quality of Fucaishu18 was a small change under the different growing patterns (Fig. 1). Among them, the latter three samples (Baisheng-H, Fucaishu18-S, and Fucaishu18-H) have similar taste values. Therefore, Fucaishu18 and Baisheng were chosen and used for the analysis of the metabolites related to taste traits by using non-targeted metabolomics.

**Fig. 1.**
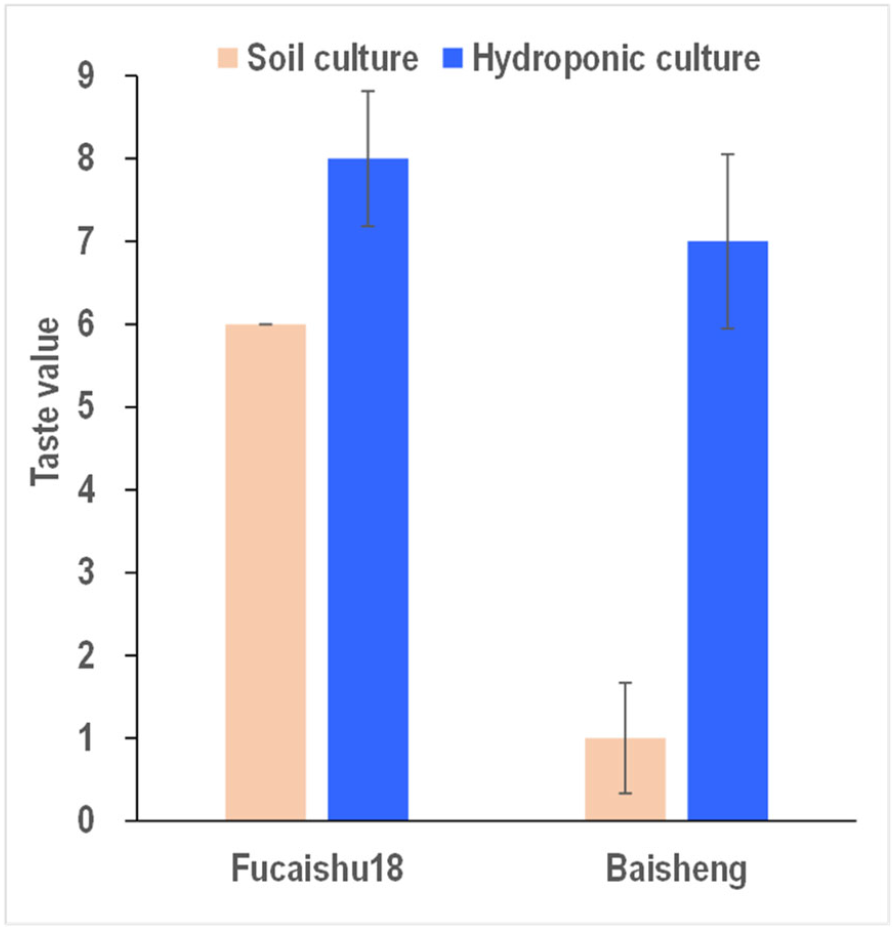
Taste values of leaves in Baisheng and Fucaishu18 under different growing patterns. Metabolite profiling of leaves in sweet potato.

By subjecting the leaves of sweet potato growth in soil and hydroponic cultures produced in the plastic house to GC-TOF-MS analysis, 518 peaks were detected and 200 metabolites could be identified by searching against LECO-Fiehn Rtx5 database (Fig. 2A). In this experiment, 5 quality control samples (QC) were used for ensuring data re-liability, and the result showed high repeatability (Fig. S1, S2) and concordance (Fig. S3). The 200 metabolites could be assigned into 7 super pathways (Fig. 2B) and sub-sequent 30 sub-pathways (Table S2) according to Kyoto Encyclopedia of Genes and Genomes (KEGG). These metabolites covered mainly the central metabolism pathways and partial secondary metabolism pathways, including 45 amino acids, 94 carbohydrates, 28 lipids, 5 CPGECs (cofactors, prosthetic groups, electron carriers), 5 nucleotides, 22 secondary metabolites, and 1 phytohormone.

**Fig. 2.**
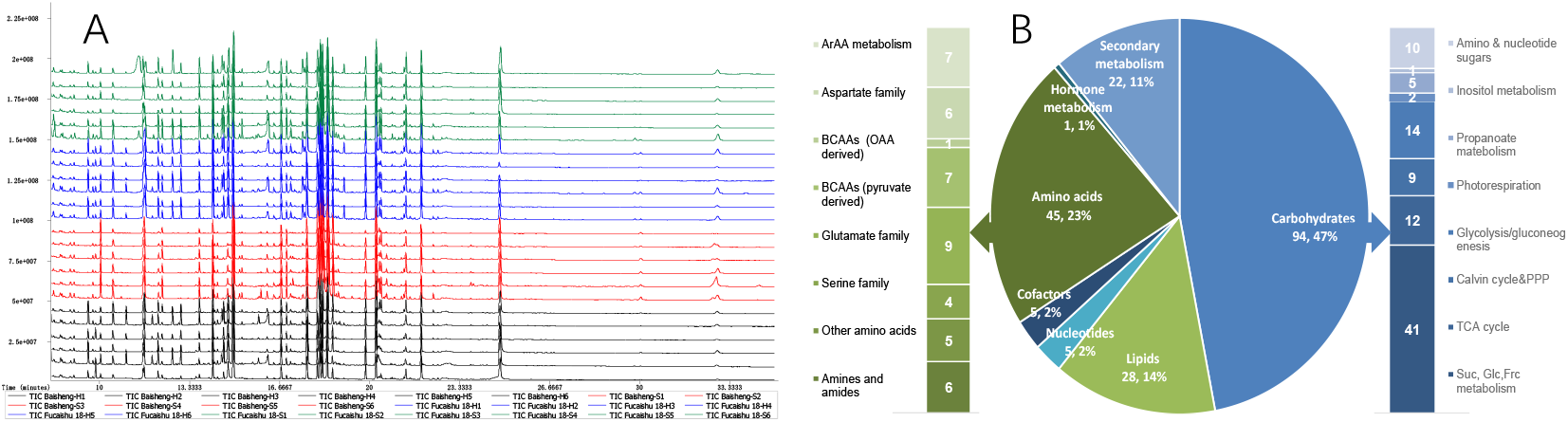
GC-TOF-MS analysis and classification of the identified metabolites in sweet potato leaves. **A**, Total Ions Chromatograph (TIC) of 4 samples with 6 replicates in sweet potato leaves via GC-TOF-MS. Green and blue graphs indicate the samples of Fucaishu18-S (soil culture) and Fucaishu18-H (hydroponic culture), respectively. red and black graphs indicate the samples of Baisheng-S (soil culture) and Baisheng-H (hydroponic culture), respectively. **B**, Distribution of identified 200 metabolites in pie chart, sub-classifications of amino acids and carbohydrates are shown with histograms.

A non-supervised principal component analysis (PCA) was performed with all 200 metabolites. As shown in Fig. 3, there was a clear separation among the 4 samples, indicating that the metabolome of Baisheng growth under soil culture (Baisheng-S, that was poor taste) was distinct from that of the other 3 samples including Baisheng-H, Fucaishu18-S, and Fucaishu18-H. Meanwhile, a heat map display of all data showed a difference cluster mode in relative abundances of Baisheng-S among the 4 samples (Fig. 4). This is the expected pattern as the result both of PCA and cluster analysis is consistent with the change in taste value between soil and hydroponic cultures of two cultivars (Fig. 1). Note that in the complexity of metabolite changes among the 4 samples, we chose 71 metabolites that the abundance was different at Baisheng-S com-pare with the other 3 samples at the same abundance, for future pathway analysis.

**Fig. 3.**
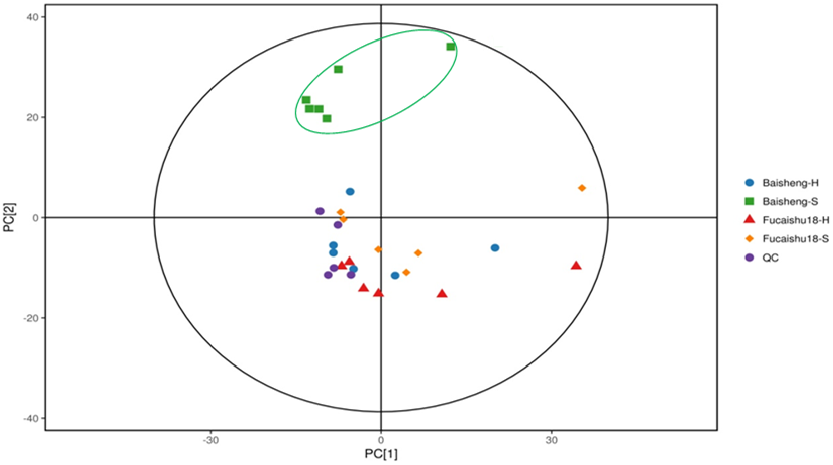
Score scatter plot of PCA model for metabolites (peaks) in leaves of Fucaishu18 and Baisheng under different growing patterns. **H**, hydroponic culture; **S**, soil culture; **QC**, quality control.

**Fig. 4.**
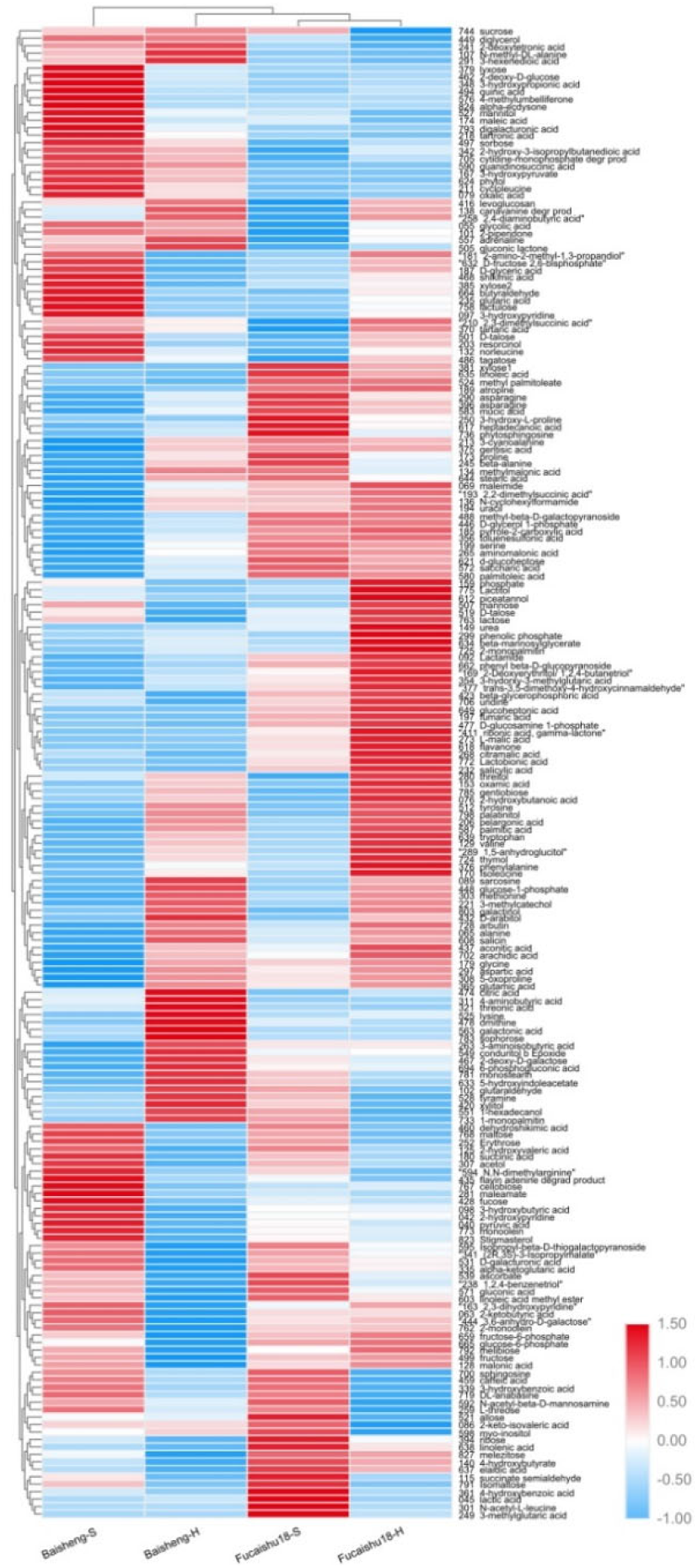
Heat map presentation of the variation of the 200 metabolites in leaves of Baisheng and Fucaishu18 under different growing patterns. **H**, hydroponic culture; **S**, soil culture. Each metabolite abundances in 4 samples were normalized with Z-score analysis.

### Metabolites and their pathway related to taste quality in sweet potato leaves

#### Reduced amino acids abundance in Baisheng-S with poor taste

Amino acid metabolism is a major part of metabolic networks in biological systems; meanwhile, the amino acids and their derivatized into numerous secondary metabolites, which are the bioactive or flavor substance in food (Kays and Wang, 2000; Lindsay, 2017). In this study, we detected 18 metabolites involve in amino acid metabolism. Except for N, N-dimethylarginine (ADMA), 3-cyanoalanine, N-amidino-L-aspartate, cycloleucine, the Baisheng-S contained a lower abundance of other 14 amino acids and their derivatives. All of four umami amino acids including Asp, Glu, Gly, and Ala were lower abundance in Baisheng-S, especially (Fig. 5, Table S2).

**Fig. 5.**
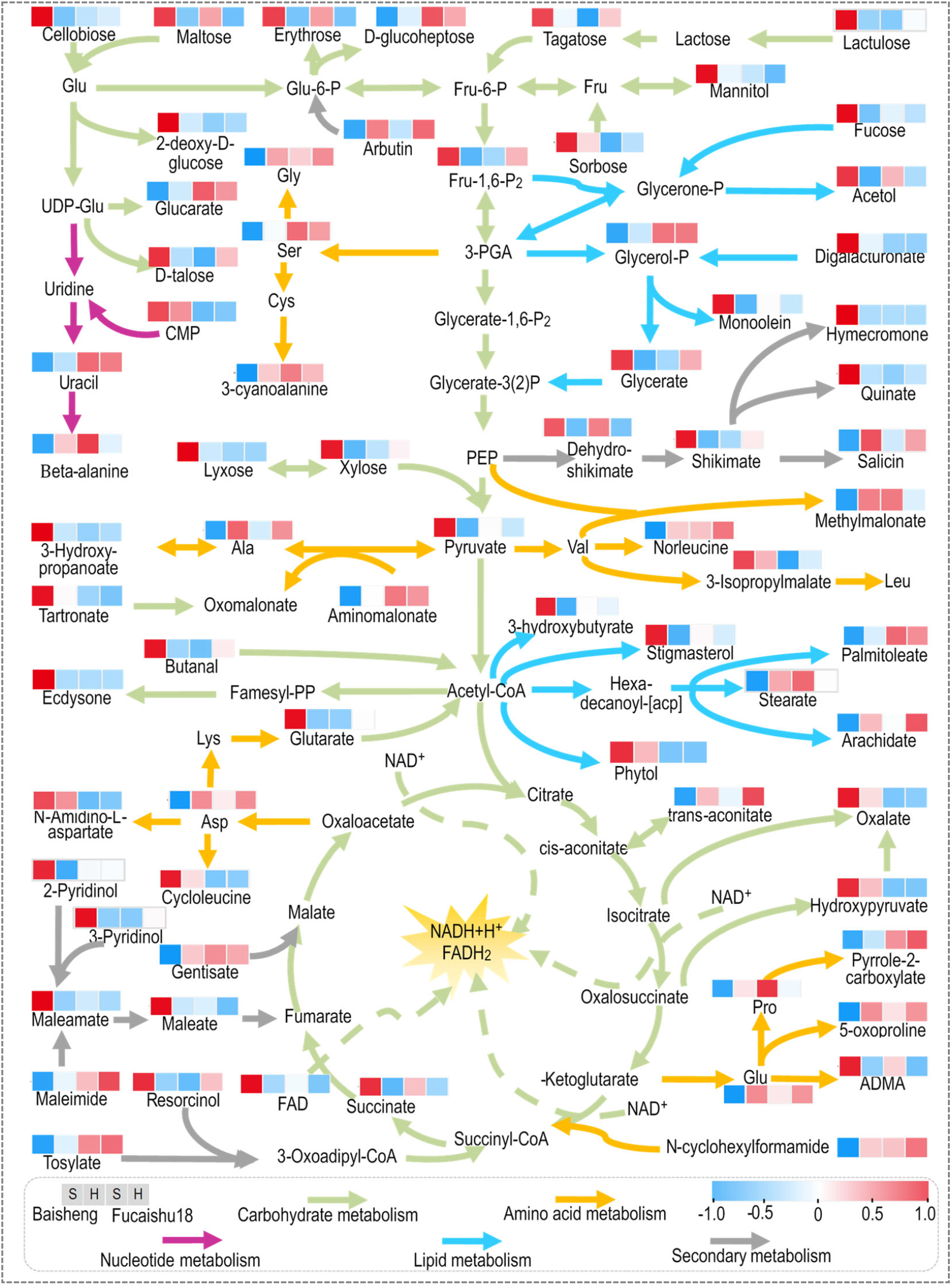
Metabolite abundances in leaves of Baisheng and Fucaishu18 under different growing patterns. The four squares under the names of metabolites indicate abundance change of Baisheng-Soil culture(S), Baisheng-Hydroponic culture(H), Fuchaishu18-Soil culture(S), Fuchaishu18-Hydroponic culture(H), respectively. Metabolite abundances of 4 samples were normalized with Z-score analysis. A red square indicates high abundance, whereas a blue square indicates low abundance.

#### Enhanced lipids synthesis, but not for ester synthesis

Plants lipids are important components in cell membrane, signal molecules, and stored energy. In this study, 10 metabolites related to lipids metabolism, that including derived from acetyl-CoA and 3-PGA. Seven of them were increased abundance in Baisheng-S, but three esters including stearate, arachidate, and palmitoleate were de-creased (Fig. 5, Table S2).

#### Increased secondary metabolism and accumulate

Plant secondary metabolites, also referred to as natural products or specialized metabolites, constitute an enormously rich reservoir of chemical biodiversity. They are provided most of the favor substances and bioactive metabolites for human health in vegetables. Thirteen secondary metabolites were detected change in Baisheng-S, among them, 9 metabolites were increased. Notably, the most important pathway for secondary metabolism, the shikimate metabolic pathway was enhanced (Fig. 5, Table S2). This suggests that secondary metabolites were accumulated and associated with worse taste.

## DISCUSSION

Metabolomics is a powerful tool for studying the quality traits of crops (Liu et al., 2019; Zhu et al., 2018; Zou et al., 2020). In previous studies, metabolomics has been used successfully for quality study in sweet potato, including anthocyanin accumulation in tuberous roots, metabolic mechanism of flesh color difference, metabolic change during postharvest storage, and metabolic diversity in leaves and roots (Wang et al., 2018; Drapal et al., 2019; He et al., 2020; Ren et al., 2021). However, the taste quality of sweet potato leaves has not been investigated so far. In this study, we used GC– QToF–MS-based untargeted metabolomic to analyze taste differences under the soil and hydroponic cultures of sweet potato leaves. A total of 200 metabolites were identified, and 71 metabolites of which were related to flavor formation in sweet potato leaves. Thus, this study provides a metabolic reference of taste quality in leafy sweet potato.

Metabolites are essential components in plants, and they can be classified as carbohydrates, organic and amino acids, vitamins, hormones, flavonoids, phenolics, and glucosinolates (Hounsome et al., 2008; Mabuchi et al., 2019). Some of these metabolites can determine the taste of leafy vegetables. Free amino acids can provide a variety of taste characteristics in leafy vegetables, such as glycine and alanine are sweetness, valine, tryptophan, and leucine are bitterness, and aspartic acid and glutamate have a sour taste (Ardö, 2006; Hounsome et al., 2008; Mabuchi et al., 2019; Toko, 2000). Another major group of sub-stances that determine taste is secondary metabolites, including flavonoids, phenolics, and alkaloids. They are the main source of bitterness and astringency (Mabuchi et al., 2019; Peterson and Dwyer, 1998; Streit et al., 2007). Our results in agreement with these above studies. Thus, we speculate that, the reason for the poor taste of sweet potato leaves under soil culture is the lower levels of metabolites regarding amino acids metabolism, whereas was accompanied by high levels of metabolites in carbohydrates and secondary metabolism. Therefore, further research should focus on the dynamic metabolic characteristics of amino acids and secondary metabolites, to explore the formation mechanism of the taste quality in leafy sweet potato, and the agronomic measures to improve the taste quality.

## MATERIALS AND METHODS

### Plant Growth and Sample Collection

The experiments were performed at the Pudang agricultural experiment farm in Fujian Academy of Agricultural Sciences (26°07′59″ N, 119°20′06″ E), Fuzhou City, China. To explore the metabolic mechanisms of taste quality of sweet potato leaves, two sweet potato cultivars including Fucaishu18 and Baisheng was used as experimental material with two types of growing patterns including hydroponic culture and soil culture under the plastic house.

Hydroponic culture (H). Eight plants in a container (40cm×28cm×15cm) of two cul-tivars were growing in half-strength Hoagland solution couple with suitable micro-nutrients (Hoagland and Arnon; 1950), and pumping air for 5 minutes every two hours in the nutrient solution.

Soil culture (S). A pot experiment was conducted using plastic pots, 15 cm in height and 12 cm in diameter. Each pot was filled with 1.5 kg clay soil, containing 0.83 g kg^−1^ total N, 10.72 mg kg^−1^ available P, and 69.15 mg kg−1 exchangeable K. For one pot, five plants of sweet potatoes were hand transplanted. The basal fertilization before trans-planting used 0.20 g N, 0.24 g P_2_O_5_, and 0.9 g K_2_O per pot.

At the 30 days after transplanted, the last leaves with fully expanded form five plants were collected from Fucaishu18 and Baisheng under different growing patterns. The four samples were named Fucaishu18-S (soil culture), Fucaishu18-H (hydroponic culture), Baisheng-S (soil culture), and Baisheng-H (hydroponic culture), respectively.

Leaves were frozen by liquid nitrogen and then stored at −80°C until analysis. Each sample was measured with six biological replicates.

### Sensory Analysis

According to the practice method from the experienced breeder of leafy sweet potato, a sensory panel of 10 trained panelists was selected in the evaluation of sensory proper-ties, with the methodology modified by Ona et al (1991) and Ishiguro et al (2004). Samples of sweet potato leaves were cooked for 2 minutes by boiling water, then evaluated imme-diately with four aspects including flavor, sweetness, bitterness, and crispiness. The leaves of Fuchaishu18 growth in soil were used as the standard with defining its evalu-ation score as 6, and other samples have a score range of 0-9 from the worst to the best by sensory of each panel.

### Metabolomic Analysis

#### Metabolites Extraction and GC-MS Analysis

Metabolite profiling was performed by Shanghai Biotree Biotech Co., Ltd. in China, according to the method of Wu *et al*. (2019). Briefly, take 60mg sample into the 2mL EP tubes, extracted with 0.48mL extraction liquid (VMethanol: VH2O = 3:1), add 20μL of adonitol (0.5mg/mL stock in dH2O) as internal standard, vortex mixing for 30s; Homogenized in ball mill for 4min at 45Hz, then ultrasound treated for 5min (incubated in ice water); Centrifuge for 15minat 12000rpm, 4°C; Transfer the supernatant (0.35mL) into a fresh 2mL GC/MS glass vial, take 40μL from each sample and pooling as QC sample. Dry completely in a vacuum concentrator without heating; Add 80μL Methoxyamination hydrochloride (20mg/mL in pyridine) incubated for 30min at 80°C; Add 100μL of the BSTFA regent (1% TMCS, v/v) to the sample aliquots, incubated for 1.5h at 70°C; Add 10μL FAMEs (Standard mixture of fatty acid methyl esters, C8-C16:1mg/mL; C18-C24:0.5mg/mL in chloroform) to the QC sample when cooling to the room temper-ature; All samples were analyzed by gas chromatography system (Agilent 7890) coupled with a Pegasus 4D time-of-flight mass spectrometer (GC-TOF-MS).

#### Data Analysis and Annotation

Chroma TOF 4.3X software of LECO Corporation and LECO-Fiehn Rtx5 database were used for raw peaks exacting, the data baselines filtering and calibration of the baseline, peak alignment, deconvolution analysis, peak identification, and integration of the peak area (Kind et al., 2009). Both mass spectrum match and retention index match were considered in metabolites identification. Remove peaks detected in <50% of QC samples or RSD>30% in QC samples (Dunn et al., 2011). Besides, internal standard normalization method was employed in this data analysis. The resulted three-dimensional data involving the peak number, sample name, and normalized peak area were fed to SIMCA software package (V14.1, MKS Data Analytics Solutions, Umea, Sweden) for principal component analysis (PCA) and orthogonal projections to latent structures-discriminate analysis (OPLS-DA). Public databases including KEGG (http://www.genome.jp/kegg/) and MetaboAnalyst (http://www.metaboanalyst.ca/) was utilized to search for the pathways of metabolites.

## Competing interests

The authors declare no competing or financial interests.

## Author contributions

Methodology, Z.L. (Zhaomiao Lin) and G.L.; software, G.L.; investigation, G.L., G.X. and Y.X.; resources, Z.L. (Zhonghua Liu), R.J., W.L. and G.X.; writing-review and editing, Z.L. (Zhaomiao Lin), G.L., H.Z., G.X., and S.Q.; visualization, Z.L. (Zhaomiao Lin); supervision, Y.Q., S.Q., and H.T.; funding acquisition, S.Q. and H.T. All authors have read and agreed to the published version of the manuscript.

## Funding

This work was supported by the Fujian Provincial Public Research Institute of Fundamental Research (2017R1026-5), the Special Fund for the Industrial Technology System Construction of Modern Agriculture of China (CARS-10-B14), the Fujian Provincial Department of Science & Technology (2017NZ0002-2), and the Fujian Academy of Agricultural Sciences Research Project (AGP2018-12).

## Supplementary information

**Fig. S1**: Total Ions Chromatograph (TIC) of 5 Quality Control Samples, 5 Internal Standards in QC samples, and Blank sample; **Fig. S2**: Principal Component Analysis plot of the peaks in 5 QC samples and 24 test samples; **Fig. S3**: High concordance among the peaks in five QC samples (r2≥0.941). **Fig. S4**: Score scatter plot of OPLS-DA model for metabolites (peaks) in leaves of Fucaishu18 and Baisheng under different growing patterns; **Fig. S5**: Pathway analysis for different metabolites in leaves of Baisheng and Fucaishu18 under different growing patterns. **Table S1**: Taste values of leaves in eight cultivars under different growing patterns; **Table S2**: Heat map of metabolite abundance in leaves of sweet potato.

## Supplementary Data

**Table S1.** Taste values of sweet potato leaves in eight cultivars under different growing patterns.

| Cultivars | Fucaishu18 | Fushu7-6 | Tainong71 | Baisheng | Fushu24 | Guangshu87 | Jinshan57 | Fushu604 |
| --- | --- | --- | --- | --- | --- | --- | --- | --- |
| Soil cultures (S) | 6.00 | 5.50±1.08 | 5.00±0.94 | 1.00±0.67 | 1.50±0.85 | 1.00±0.67 | 1.50±0.97 | 4.50±1.27 |
| Hydroponics (H) | 8.00±0.82 | 7.50±0.71 | 8.00±1.25 | 7.00±1.05 | 4.50±1.90 | 4.00±0.82 | 5.00±1.41 | 7.00±0.94 |
| Differences | 2.00 | 2.00 | 3.00 | 6.00 | 3.00 | 3.00 | 3.50 | 2.50 |

Fucaishu18, Fushu7-6, Tainong71 are leafy sweetpotatoes for leaf vegetable, while other cultivars are common sweetpotatoes for tuber production.
The leaves of Fuchaishu18 growth in soil were used as the standard with defining its evaluation score as 6.

**Fig. S1.**
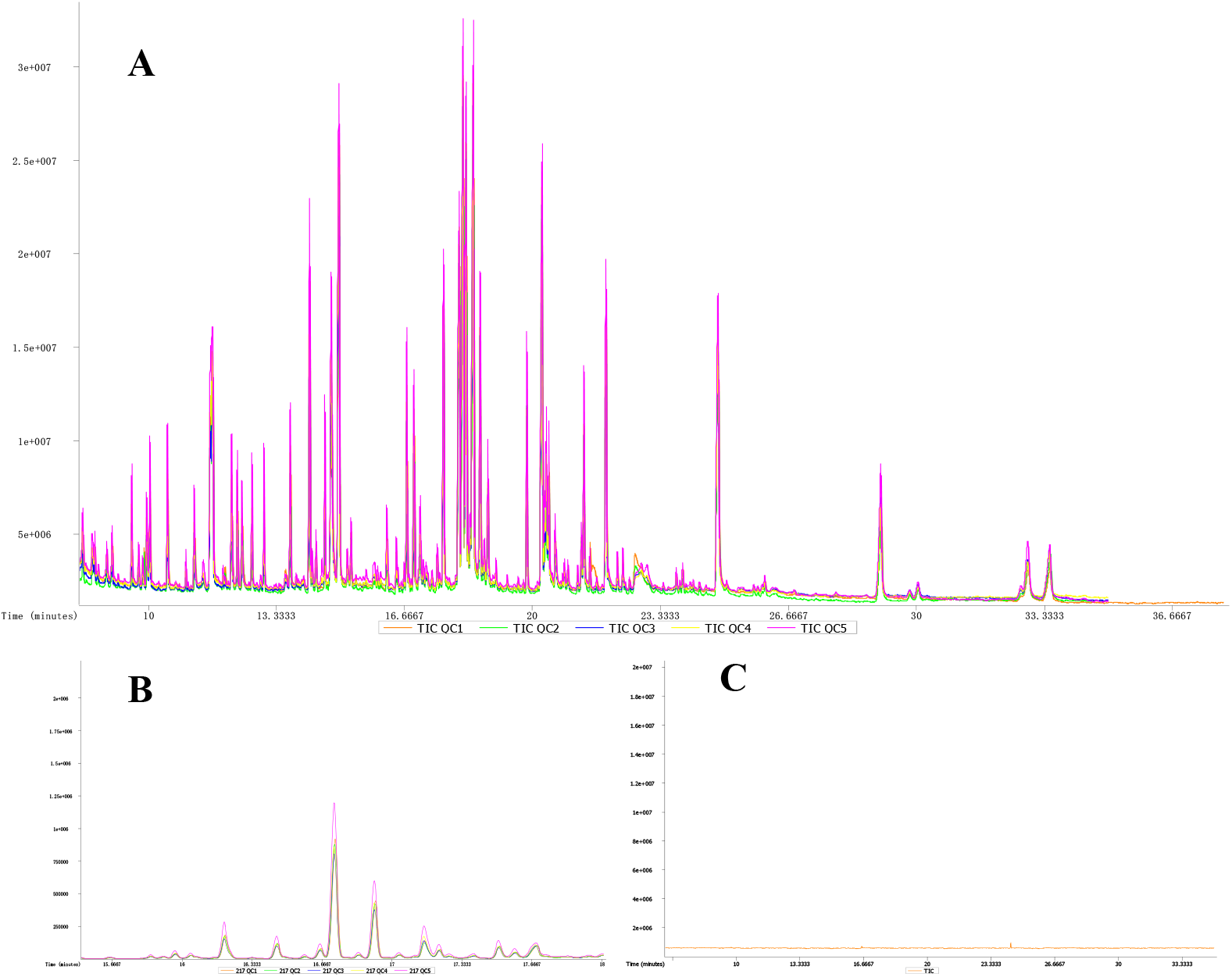
Total Ions Chromatograph (TIC) of 5 Quality Control Samples (A), 5 Internal Standards in QC samples (B), and Blank sample (C).

**Fig. S2.**
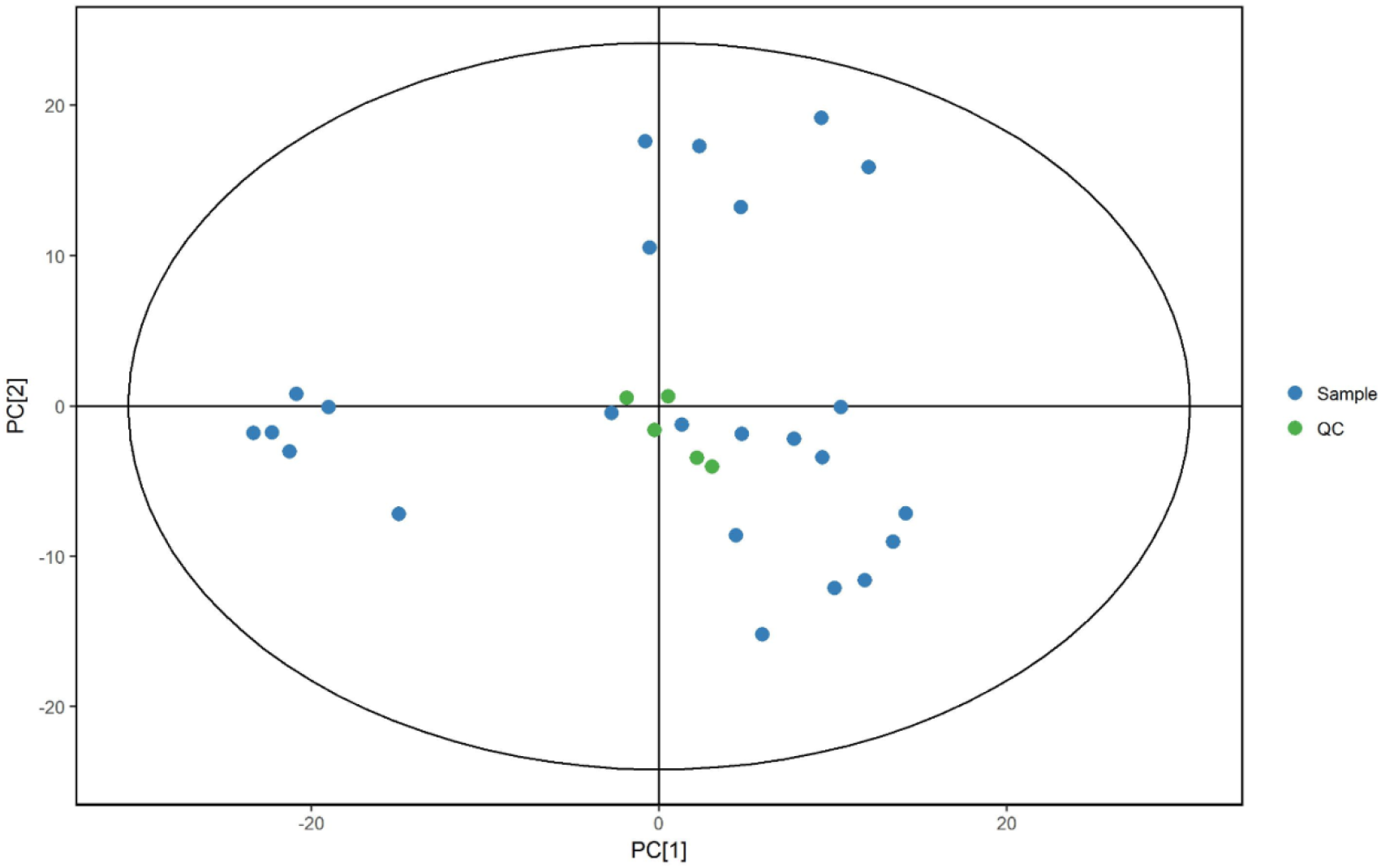
Principal Component Analysis plot of the peaks in 5 QC samples and 24 test samples. Green spots indicate the QC samples, while blue spots indicate the test samples.

**Fig. S3.**
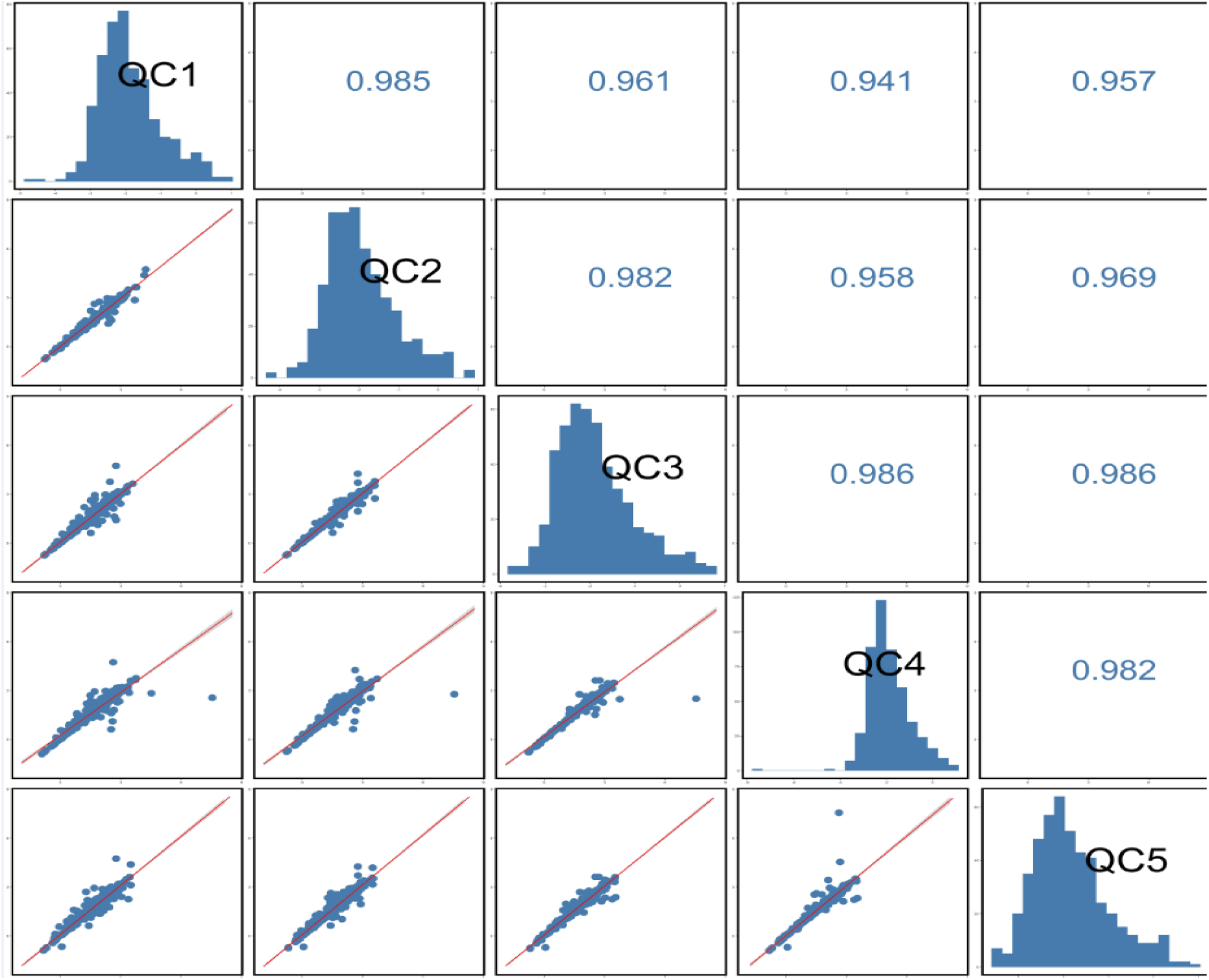
High concordance among the peaks in five QC samples (r^2^≥0.941).

**Fig. S4.**
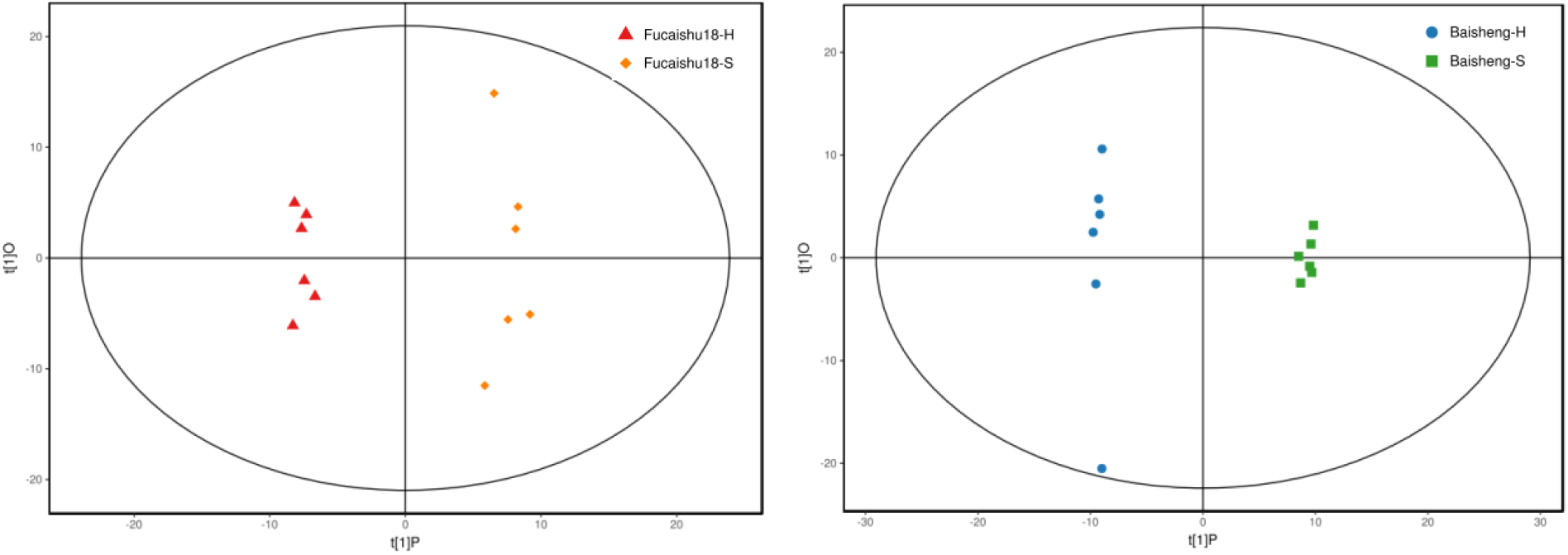
Score scatter plot of OPLS-DA model for metabolites (peaks) in leaves of Fucaishu18(A) and Baisheng(B) under different growing patterns.

**Fig. S5.**
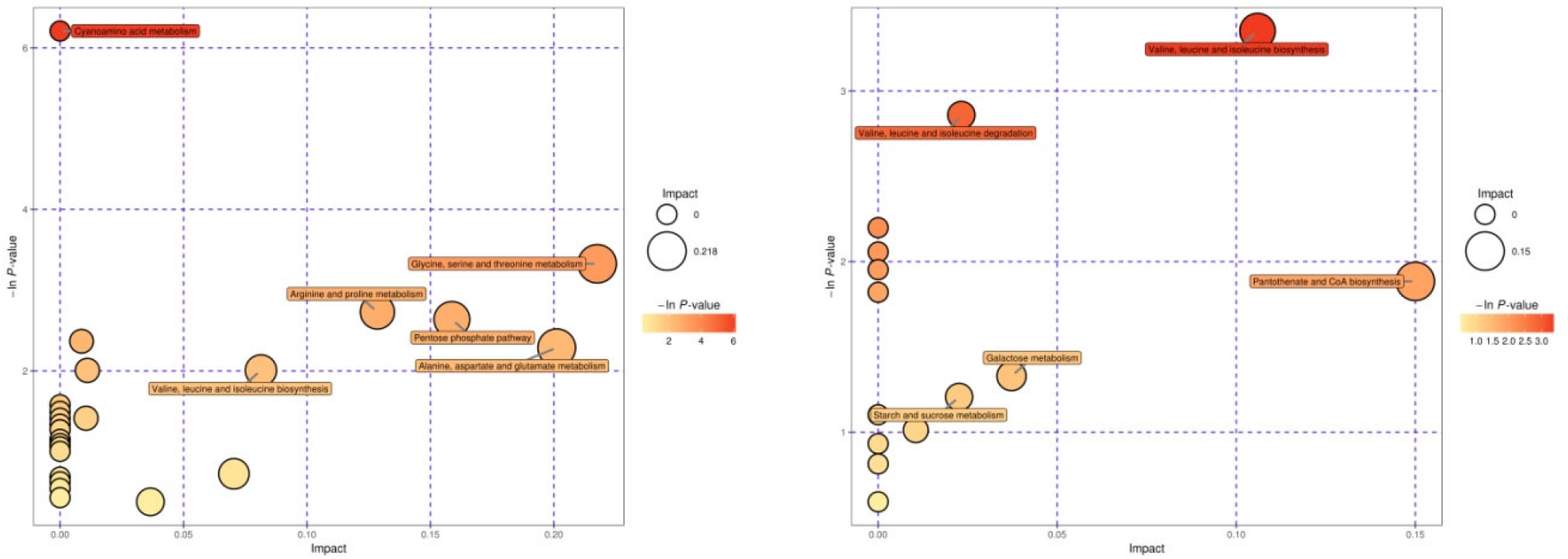
Pathway analysis for different metabolites in leaves of Baisheng (A) and Fucaishu18 (B) under different growing patterns.

## Notes

### Competing Interest Statement

The authors have declared no competing interest.

