## Supplemental Table S2 for "Metabolic characteristics of taste differences under the soil and hydroponic cultures of sweet potato leaves by using non-targeted metabolomics"

### Table S2. Heat map of metabolite abundance in leaves of sweetpotato.

At the column of Relative Contents, a light green cells indicate low abundance of this metabolite of all metabolites, whereas a dark green cells indicate high abundance. At the column of Normalization Valuce, a red shaded cells indicate that this metabolite is high abundance among four samples, whereas a blue shaded cells indicate low abundance.

**S,** Soil culture; **H,** Hydroponic culture**.**

| ID | Super Pathway | Sub Pathway | biochemical Name | CAS | Relative Contents | | | | Normalization Valuce of Relative Contents in each row among four samples by Z-scrose analysis | | | |
|---|---|---|---|---|---|---|---|---|---|---|---|---|
| | | | | | Baisheng-S | Baisheng-H | Fucaishu18-S | Fucaishu18-H | Baisheng-S | Baisheng-H | Fucaishu18-S | Fucaishu18-H |
| 376 | | | phenylalanine | 63-91-2 | 0.0000 | 0.0159 | 0.0096 | 0.0416 | -0.9419 | -0.0486 | -0.4047 | 1.3952 |
| 460 | | | dehydroshikimate | 2922-42-1 | 0.0002 | 0.0001 | 0.0002 | 0.0001 | 0.9777 | -0.8958 | 0.7459 | -0.8278 |
| 468 | | Aromatic amino acid metabolism (PEP derived) | shikimate | 138-59-0 | 0.2910 | 0.1733 | 0.1940 | 0.2279 | 1.3488 | -0.9375 | -0.5353 | 0.1240 |
| 512 | | | tyrosine | 60-18-4 | 0.0000 | 0.0245 | 0.0011 | 0.0302 | -0.8924 | 0.6727 | -0.8194 | 1.0391 |
| 528 | | | tyramine | 51-67-2 | 0.0058 | 0.0171 | 0.0120 | 0.0029 | -0.5743 | 1.2064 | 0.3964 | -1.0284 |
| 557 | | | adrenaline | 51-43-4 | 0.0010 | 0.0012 | 0.0006 | 0.0009 | 0.2943 | 1.0698 | -1.3296 | -0.0345 |
| 639 | | | tryptophan | 73-22-3 | 0.0000 | 0.1951 | 0.0591 | 0.3177 | -1.0047 | 0.3665 | -0.5898 | 1.2281 |
| 132 | | | norleucine | 327-57-1 | 0.0221 | 0.0191 | 0.0174 | 0.0196 | 1.3087 | -0.2387 | -1.1085 | 0.0385 |
| 290 | | | asparagine1 | 70-47-3 | 0.0000 | 0.3324 | 0.8192 | 0.4541 | -1.1868 | -0.2040 | 1.2352 | 0.1556 |
| 297 | | Aspartate family (OAA derived) | aspartate | 56-84-8 | 0.0491 | 1.3765 | 1.0240 | 1.3692 | -1.4472 | 0.6741 | 0.1107 | 0.6624 |
| 303 | | | methionine | 63-68-3 | 0.0000 | 0.0908 | 0.0302 | 0.0779 | -1.1793 | 0.9731 | -0.4631 | 0.6692 |
| 396 | | | asparagine2 | 70-47-3 | 0.0000 | 0.2053 | 0.5408 | 0.3575 | -1.2024 | -0.3078 | 1.1545 | 0.3557 |
| 525 | | | lysine | 56-87-1 | 0.0146 | 0.0497 | 0.0195 | 0.0241 | -0.7893 | 1.4539 | -0.4778 | -0.1869 |
| 170 | | Branched Chain Amir | Isoleucine | 73-32-5 | 0.0000 | 0.1514 | 0.0484 | 0.3421 | -0.8941 | 0.1052 | -0.5746 | 1.3636 |
| 65 | | | alanine | 56-41-7 | 0.0984 | 3.5915 | 1.7405 | 3.1672 | -1.2981 | 0.9127 | -0.2588 | 0.6442 |
| 86 | | | 2-keto-isovalerate | 759-05-7 | 0.0026 | 0.0026 | 0.0033 | 0.0009 | 0.2674 | 0.2284 | 0.9254 | -1.4212 |
| 107 | | Branched Chain Amino Acids (pyruvate derived) | N-methyl-DL-alanine | 600-21-5 | 0.0073 | 0.0076 | 0.0069 | 0.0068 | 0.3911 | 1.2295 | -0.7048 | -0.9158 |
| 129 | | | valine | 72-18-4 | 0.0051 | 0.3415 | 0.1251 | 0.5530 | -1.0378 | 0.3526 | -0.5417 | 1.2269 |
| 211 | | | cycloleucine | 52-52-8 | 0.0126 | 0.0089 | 0.0053 | 0.0062 | 1.3257 | 0.2008 | -0.8993 | -0.6272 |
| 301 | | | N-acetyl-L-leucine | 1188-21-2 | 0.0000 | 0.0000 | 0.0070 | 0.0000 | -0.5000 | -0.5000 | 1.5000 | -0.5000 |
| 341 | | | (2R,3S)-3-Isopropylmalate | 921-28-8 | 0.2298 | 0.0884 | 0.2102 | 0.1713 | 0.8772 | -1.3819 | 0.5634 | -0.0587 |
| 115 | | | succinate semialdehyde | 692-29-5 | 0.0016 | 0.0007 | 0.0026 | 0.0010 | 0.1338 | -0.9267 | 1.3475 | -0.5546 |
| 138 | Amino acids | | canavanine degr prod | 543-38-4 | 0.0135 | 0.0151 | 0.0121 | 0.0144 | -0.2275 | 1.0150 | -1.2904 | 0.5029 |
| 173 | | | proline | 147-85-3 | 0.0000 | 0.0997 | 0.1707 | 0.0830 | -1.2604 | 0.1617 | 1.1746 | -0.0759 |
| 250 | | Glutamate family (alpha-ketoglutarate derived) | 3-hydroxy-L-proline | 119677-21-3 | 0.0011 | 0.0011 | 0.0084 | 0.0037 | -0.7101 | -0.7219 | 1.4050 | 0.0270 |
| 308 | | | 5-oxoproline | 98-79-3 | 1.3547 | 5.0033 | 4.1698 | 4.9585 | -1.4624 | 0.6576 | 0.1733 | 0.6315 |
| 311 | | | 4-aminobutyrate | 1956-12-2 | 0.1331 | 0.3491 | 0.0697 | 0.0775 | -0.1853 | 1.4647 | -0.6697 | -0.6097 |
| 365 | | | glutamate | 56-86-0 | 0.0000 | 0.0453 | 0.0339 | 0.0442 | -1.4551 | 0.6811 | 0.1427 | 0.6312 |
| 478 | | | ornithine | 70-26-8 | 0.0016 | 0.0518 | 0.0141 | 0.0079 | -0.7656 | 1.4610 | -0.2108 | -0.4847 |
| 594 | | | N,N-dimethylarginine (ADMA) | 30315-93-6 | 0.0015 | 0.0003 | 0.0009 | 0.0001 | 1.3024 | -0.6834 | 0.2585 | -0.8775 |
| 63 | | | 2-ketobutyrate | 600-18-0 | 0.0119 | 0.0027 | 0.0090 | 0.0090 | 0.9682 | -1.4029 | 0.2192 | 0.2154 |
| 89 | | Serine family (phosphoglycerate derived) | sarcosine | 107-97-1 | 0.0021 | 0.0101 | 0.0035 | 0.0075 | -1.0129 | 1.1678 | -0.6244 | 0.4694 |
| 179 | | | glycine | 56-40-6 | 0.1623 | 1.6635 | 1.4785 | 1.7718 | -1.4802 | 0.5277 | 0.2801 | 0.6724 |
| 199 | | | serine | 56-45-1 | 0.0026 | 0.4590 | 0.7701 | 0.6710 | -1.3872 | -0.0489 | 0.8633 | 0.5727 |
| 213 | | | 3-cyanoalanine | 6232-19-5 | 0.0001 | 0.0820 | 0.1030 | 0.0862 | -1.4706 | 0.3072 | 0.7636 | 0.3999 |
| 258 | | | 2,4-diaminobutyrate | 305-62-4 | 0.0051 | 0.0065 | 0.0040 | 0.0061 | -0.2797 | 0.9576 | -1.2810 | 0.6030 |

| | | | | | | | | | | | |
|---|---|---|---|---|---|---|---|---|---|---|---|
| 263 | Other amino acids | 3-aminoisobutyrate | 144-90-1 | 0.0070 | 0.0085 | 0.0079 | 0.0079 | -1.3532 | 1.0612 | 0.1524 | 0.1395 |
| 265 | | aminomalonate | 1068-84-4 | 0.0046 | 0.0074 | 0.0090 | 0.0086 | -1.4100 | -0.0010 | 0.7964 | 0.6146 |
| 590 | | N-amidino-L-aspartate | 6133-30-8 | 0.0006 | 0.0005 | 0.0001 | 0.0002 | 1.0606 | 0.6429 | -0.9250 | -0.7785 |
| 69 | Amines and amides | maleimide | 541-59-3 | 0.0097 | 0.0243 | 0.0307 | 0.0388 | -1.3148 | -0.1263 | 0.3913 | 1.0498 |
| 92 | | Lactamide | 2043-43-8 | 0.0003 | 0.0015 | 0.0031 | 0.0049 | -1.0821 | -0.4659 | 0.3191 | 1.2289 |
| 136 | | N-cyclohexylformamide | 766-93-8 | 0.0010 | 0.0018 | 0.0018 | 0.0020 | -1.4607 | 0.2992 | 0.3546 | 0.8069 |
| 149 | | urea | 57-13-6 | 0.0000 | 0.0100 | 0.0253 | 0.0969 | -0.7545 | -0.5262 | -0.1763 | 1.4570 |
| 281 | | maleamate | 557-24-4 | 0.0074 | 0.0002 | 0.0015 | 0.0006 | 1.4825 | -0.6576 | -0.2907 | -0.5341 |
| 592 | | N-acetyl-beta-D-mannosamine | 7772-94-3 | 0.0027 | 0.0025 | 0.0029 | 0.0021 | 0.5495 | -0.2023 | 0.9659 | -1.3131 |
| 289 | Sucrose, glucose,fructose metabolism | 1,5-anhydroglucitol | 154-58-5 | 0.0003 | 0.0006 | 0.0004 | 0.0008 | -0.9881 | 0.2512 | -0.5506 | 1.2874 |
| 416 | | levoglucosan | 498-07-7 | 0.0188 | 0.0209 | 0.0131 | 0.0196 | 0.2061 | 0.8129 | -1.4522 | 0.4333 |
| 428 | | fucose | 2438-80-4 | 0.0070 | 0.0061 | 0.0064 | 0.0063 | 1.4387 | -0.8515 | -0.1660 | -0.4212 |
| 444 | | 3,6-anhydro-D-galactose | 14122-18-0 | 0.0142 | 0.0078 | 0.0122 | 0.0128 | 0.8932 | -1.4282 | 0.1690 | 0.3660 |
| 462 | | 2-deoxy-D-glucose | 154-17-6 | 0.0424 | 0.0248 | 0.0209 | 0.0224 | 1.4805 | -0.2836 | -0.6740 | -0.5228 |
| 467 | | 2-deoxy-D-galactose | 1949-89-9 | 0.0262 | 0.1142 | 0.0741 | 0.0596 | -1.1612 | 1.2524 | 0.1533 | -0.2444 |
| 486 | | tagatose | 87-81-0 | 0.0031 | 0.0023 | 0.0016 | 0.0026 | 1.0893 | -0.0932 | -1.3069 | 0.3108 |
| 488 | | methyl-beta-D-galactopyranosid | 1824-94-8 | 0.0016 | 0.0280 | 0.0716 | 0.0676 | -1.2131 | -0.4247 | 0.8778 | 0.7600 |
| 499 | | fructose | 57-48-7 | 5.4100 | 3.3019 | 5.2143 | 5.4811 | 0.5370 | -1.4911 | 0.3487 | 0.6054 |
| 501 | | D-talose1 | 2595-98-4 | 0.0344 | 0.0200 | 0.0144 | 0.0271 | 1.2012 | -0.4561 | -1.1045 | 0.3594 |
| 505 | | gluconic lactone | 90-80-2 | 0.0060 | 0.0068 | 0.0043 | 0.0050 | 0.4520 | 1.1404 | -1.1333 | -0.4590 |
| 507 | | mannose | 3458-28-4 | 2.2542 | 1.3667 | 1.6067 | 2.5943 | 0.5265 | -1.0377 | -0.6148 | 1.1260 |
| 519 | | D-talose2 | 2595-98-4 | 0.5310 | 0.4082 | 0.5041 | 0.6322 | 0.1318 | -1.2015 | -0.1604 | 1.2301 |
| 521 | | allose | 2595-97-3 | 0.0164 | 0.0153 | 0.0274 | 0.0078 | -0.0374 | -0.1772 | 1.3206 | -1.1060 |
| 527 | | mannitol | 87-78-5 | 0.0747 | 0.0650 | 0.0636 | 0.0599 | 1.4098 | -0.1244 | -0.3506 | -0.9349 |
| 531 | | D-galacturonate | 685-73-4 | 0.0084 | 0.0017 | 0.0071 | 0.0060 | 0.8862 | -1.4149 | 0.4552 | 0.0734 |
| 549 | | conduritol b Epoxide | 6090-95-5 | 0.0803 | 0.1669 | 0.1184 | 0.1221 | -1.1737 | 1.2698 | -0.0998 | 0.0037 |
| 563 | | galactonate | 576-36-3 | 0.0000 | 0.0050 | 0.0000 | 0.0000 | -0.5000 | 1.5000 | -0.5000 | -0.5000 |
| 571 | | gluconate | 526-95-4 | 0.0056 | 0.0008 | 0.0070 | 0.0042 | 0.4633 | -1.3614 | 0.9632 | -0.0650 |
| 572 | | Glucarate | 576-42-1 | 0.0039 | 0.0066 | 0.0101 | 0.0092 | -1.2775 | -0.2899 | 0.9501 | 0.6173 |
| 595 | | Isopropyl-beta-D-thiogalactopyr | 367-93-1 | 0.0312 | 0.0140 | 0.0320 | 0.0253 | 0.6723 | -1.4006 | 0.7649 | -0.0365 |
| 621 | | d-glucoheptose | 62475-58-5 | 0.0159 | 0.0352 | 0.0563 | 0.0476 | -1.3043 | -0.2030 | 1.0002 | 0.5071 |
| 632 | | D-fructose 2,6-bisphosphate | 79082-92-1 | 0.0531 | 0.0260 | 0.0319 | 0.0440 | 1.1819 | -1.0482 | -0.5646 | 0.4309 |
| 649 | | glucoheptonate | 23351-51-1 | 0.0180 | 0.0176 | 0.0251 | 0.0306 | -0.7769 | -0.8405 | 0.3718 | 1.2457 |
| 662 | | phenyl beta-D-glucopyranoside | 1464-44-4 | 0.2919 | 0.3345 | 0.4320 | 0.4961 | -1.0446 | -0.5846 | 0.4682 | 1.1610 |
| 694 | | 6-phosphogluconate | 53411-70-4 | 0.0285 | 0.0480 | 0.0385 | 0.0368 | -1.1806 | 1.2572 | 0.0678 | -0.1444 |
| 744 | | sucrose | 57-50-1 | 2.2156 | 2.5231 | 2.3622 | 0.8894 | 0.2910 | 0.7013 | 0.4866 | -1.4788 |
| 758 | | lactulose | 4618-18-2 | 0.0107 | 0.0037 | 0.0033 | 0.0057 | 1.4226 | -0.6333 | -0.7576 | -0.0317 |
| 763 | | lactose | 63-42-3 | 0.0018 | 0.0006 | 0.0014 | 0.0025 | 0.3315 | -1.2536 | -0.2072 | 1.1294 |
| 767 | | cellobiose | 528-50-7 | 0.0022 | 0.0006 | 0.0009 | 0.0010 | 1.4528 | -0.8245 | -0.3922 | -0.2361 |
| 768 | | maltose | 69-79-4 | 0.0250 | 0.0204 | 0.0236 | 0.0201 | 1.1389 | -0.7680 | 0.5372 | -0.9080 |
| 772 | | Lactobionate | 96-82-2 | 0.0006 | 0.0004 | 0.0014 | 0.0024 | -0.6733 | -0.8569 | 0.1987 | 1.3315 |
| 775 | | Lactitol | 585-86-4 | 0.0029 | 0.0023 | 0.0028 | 0.0050 | -0.2540 | -0.7957 | -0.4109 | 1.4606 |
| 783 | | sophorose | 534-46-3 | 0.0000 | 0.0034 | 0.0002 | 0.0002 | -0.5660 | 1.4983 | -0.4644 | -0.4679 |
| 785 | | gentiobiose | 554-91-6 | 0.0609 | 0.1014 | 0.0474 | 0.1317 | -0.6362 | 0.4179 | -0.9856 | 1.2038 |
| 791 | | Isomaltose | 499-40-1 | 0.0029 | 0.0000 | 0.0051 | 0.0000 | 0.3653 | -0.8076 | 1.2499 | -0.8076 |
| 792 | | melibiose | 66009-10-7 | 0.0071 | 0.0005 | 0.0053 | 0.0084 | 0.5063 | -1.3967 | -0.0001 | 0.8905 |
| 793 | | digalacturonate | 5894-59-7 | 0.0198 | 0.0061 | 0.0019 | 0.0017 | 1.4575 | -0.1518 | -0.6426 | -0.6631 |

| | | | | | | | | | | | | |
|---|---|---|---|---|---|---|---|---|---|---|---|---|
| 798 | | | palatinitol | 64519-82-0 | 0.0037 | 0.0051 | 0.0038 | 0.0054 | -0.8829 | 0.6913 | -0.8327 | 1.0243 |
| 803 | | | galactinol | 3687-64-7 | 0.0070 | 0.0154 | 0.0063 | 0.0144 | -0.7863 | 0.9684 | -0.9358 | 0.7537 |
| 827 | | | melezitose | 597-12-6 | 1.4462 | 1.1039 | 1.7023 | 1.6118 | -0.0752 | -1.3734 | 0.8961 | 0.5525 |
| 140 | | | 4-hydroxybutyrate | 502-85-2 | 0.0019 | 0.0007 | 0.0041 | 0.0032 | -0.3810 | -1.2028 | 1.0694 | 0.5144 |
| 174 | | | maleate | 110-16-7 | 0.0012 | 0.0005 | 0.0005 | 0.0003 | 1.4330 | -0.3269 | -0.2154 | -0.8906 |
| 180 | | | succinate | 110-15-6 | 0.0683 | 0.0329 | 0.0547 | 0.0397 | 1.2282 | -1.0116 | 0.3633 | -0.5798 |
| 193 | | | 2,2-dimethylsuccinate | 597-43-3 | 0.0010 | 0.0020 | 0.0022 | 0.0026 | -1.4243 | 0.1288 | 0.4072 | 0.8882 |
| 197 | | | fumarate | 110-17-8 | 0.0185 | 0.0178 | 0.0283 | 0.0341 | -0.7838 | -0.8689 | 0.4623 | 1.1903 |
| 210 | Carbohydrates | TCA cycle | 2,3-dimethylsuccinate | 13545-04-5 | 0.0632 | 0.0603 | 0.0489 | 0.0662 | 0.4696 | 0.0865 | -1.4220 | 0.8659 |
| 268 | | | citramalate | 2306-22-1 | 0.0045 | 0.0030 | 0.0061 | 0.0094 | -0.4636 | -0.9925 | 0.1198 | 1.3363 |
| 273 | | | L-malate | 97-67-6 | 0.5641 | 0.5710 | 0.7918 | 1.0548 | -0.7824 | -0.7526 | 0.2003 | 1.3347 |
| 307 | | | acetol | 116-09-6 | 0.0240 | 0.0076 | 0.0185 | 0.0119 | 1.1757 | -1.0948 | 0.4156 | -0.4964 |
| 335 | | | alpha-ketoglutarate | 328-50-7 | 0.0613 | 0.0284 | 0.0549 | 0.0495 | 0.8965 | -1.4117 | 0.4459 | 0.0693 |
| 437 | | | aconitate | 4023-65-8 | 0.0012 | 0.0032 | 0.0026 | 0.0040 | -1.3161 | 0.3888 | -0.1224 | 1.0497 |
| 474 | | | citrate | 5949-29-1 | 1.9060 | 2.5999 | 1.6734 | 1.8605 | -0.2561 | 1.4532 | -0.8290 | -0.3681 |
| 55 | | | glycolate | 79-14-1 | 0.0647 | 0.0603 | 0.0371 | 0.0526 | 0.9094 | 0.5444 | -1.3671 | -0.0867 |
| 79 | | | oxalate | 144-62-7 | 0.0037 | 0.0032 | 0.0028 | 0.0029 | 1.3366 | 0.1940 | -0.8120 | -0.7186 |
| 102 | | | glutaraldehyde | 111-30-8 | 0.0015 | 0.0020 | 0.0019 | 0.0015 | -0.8562 | 1.1360 | 0.5446 | -0.8244 |
| 167 | | Calvin cycle and pentose phosphate | 3-hydroxypyruvate | 1113-60-6 | 0.0797 | 0.0466 | 0.0012 | 0.0016 | 1.2441 | 0.3752 | -0.8157 | -0.8037 |
| 169 | | | 2-Deoxyerythritol/ 1,2,4-butanet | 3068-00-6 | 0.0088 | 0.0114 | 0.0137 | 0.0201 | -0.9770 | -0.4358 | 0.0516 | 1.3612 |
| 181 | | | 2-amino-2-methyl-1,3-propandi | 115-69-5 | 0.0066 | 0.0050 | 0.0053 | 0.0064 | 0.9295 | -1.0602 | -0.6430 | 0.7736 |
| 252 | | | Erythrose | 583-50-6 | 0.0054 | 0.0021 | 0.0045 | 0.0021 | 1.1207 | -0.8429 | 0.5664 | -0.8442 |
| 259 | | | L-threose | 95-44-3 | 0.0151 | 0.0093 | 0.0145 | 0.0041 | 0.8486 | -0.2729 | 0.7220 | -1.2977 |
| 348 | | | 3-hydroxypropionate | 503-66-2 | 0.1168 | 0.0724 | 0.0651 | 0.0690 | 1.4883 | -0.3468 | -0.6522 | -0.4893 |
| 40 | | | pyruvate | 127-17-3 | 0.0646 | 0.0226 | 0.0412 | 0.0347 | 1.3496 | -1.0294 | 0.0241 | -0.3443 |
| 128 | | | malonate | 141-82-2 | 0.0084 | 0.0056 | 0.0079 | 0.0085 | 0.5769 | -1.4692 | 0.2109 | 0.6813 |
| 218 | | | tartronate | 80-69-3 | 0.0065 | 0.0041 | 0.0029 | 0.0025 | 1.3970 | 0.0332 | -0.6039 | -0.8262 |
| 235 | | | glutarate | 110-94-1 | 0.0207 | 0.0096 | 0.0098 | 0.0133 | 1.4174 | -0.7235 | -0.6838 | -0.0102 |
| 241 | | | 2-deoxytetronate | 1518-61-2 | 0.0051 | 0.0066 | 0.0017 | 0.0001 | 0.5833 | 1.0723 | -0.5637 | -1.0919 |
| 249 | | | 3-methylglutarate | 626-51-7 | 2.0659 | 2.0662 | 2.8069 | 2.1111 | -0.5409 | -0.5399 | 1.4974 | -0.4166 |
| 339 | | gluconeogenesis/Glycolysis | 3-hydroxybenzoate | 1999-6-9 | 0.0166 | 0.0118 | 0.0168 | 0.0100 | 0.8287 | -0.5827 | 0.8627 | -1.1088 |
| 342 | | | 3-Isopropylmalate | 16048-89-8 | 0.0218 | 0.0189 | 0.0113 | 0.0158 | 1.0782 | 0.4281 | -1.2629 | -0.2435 |
| 370 | | | tartarate | 133-37-9 | 0.0010 | 0.0010 | 0.0008 | 0.0011 | 0.6416 | 0.0829 | -1.4391 | 0.7146 |
| 423 | | | beta-glycerophosphorate | 819-83-0 | 0.0051 | 0.0036 | 0.0067 | 0.0081 | -0.3927 | -1.1705 | 0.4255 | 1.1378 |
| 446 | | | D-glycerol 1-phosphate | 34363-28-5 | 0.0998 | 0.1115 | 0.1258 | 0.1255 | -1.2682 | -0.3325 | 0.8111 | 0.7895 |
| 448 | | | glucose-1-phosphate | 59-56-3 | 0.0074 | 0.0114 | 0.0081 | 0.0104 | -1.0072 | 1.0869 | -0.6727 | 0.5930 |
| 659 | | | fructose-6-phosphate | 643-13-0 | 0.0033 | 0.0000 | 0.0034 | 0.0042 | 0.3081 | -1.4668 | 0.3766 | 0.7821 |
| 665 | | | glucose-6-phosphate | 56-73-5 | 0.0035 | 0.0002 | 0.0047 | 0.0054 | 0.0347 | -1.4151 | 0.5426 | 0.8378 |
| 598 | | Inositol metabolism | myo-inositol | 87-89-8 | 2.3858 | 2.4536 | 2.6560 | 2.0556 | -0.0077 | 0.2640 | 1.0755 | -1.3318 |
| 187 | | Photorespiration | D-glycerate | 6000-40-4 | 0.4286 | 0.2570 | 0.2955 | 0.3750 | 1.1582 | -1.0608 | -0.5627 | 0.4653 |
| 497 | | | sorbose | 3615-56-3 | 9.7110 | 5.3182 | 0.0000 | 2.6167 | 1.2780 | 0.2187 | -1.0638 | -0.4328 |
| 45 | | | lactate | 50-21-5 | 0.0490 | 0.0513 | 0.0748 | 0.0477 | -0.5217 | -0.3430 | 1.4898 | -0.6250 |
| 76 | | | 2-hydroxybutanoate | 565-70-8 | 0.4003 | 0.7695 | 0.3013 | 1.1422 | -0.6605 | 0.3032 | -0.9186 | 1.2759 |
| 98 | | Propanoate matebolism | 3-hydroxybutyrate | 300-85-6 | 0.0337 | 0.0255 | 0.0294 | 0.0289 | 1.2889 | -1.1494 | -0.0073 | -0.1322 |
| 134 | | | methylmalonate | 516-05-2 | 0.0057 | 0.0066 | 0.0066 | 0.0062 | -1.3331 | 0.7476 | 0.7889 | -0.2035 |
| 291 | | | 3-hexenedioate | 4436-74-2 | 0.0043 | 0.0063 | 0.0020 | 0.0014 | 0.3664 | 1.2404 | -0.6693 | -0.9376 |
| 280 | | | threitol | 7493-90-5 | 0.1489 | 0.1633 | 0.1284 | 0.1818 | -0.2962 | 0.3412 | -1.2046 | 1.1596 |

| | | | | | | | | | | | | |
|---|---|---|---|---|---|---|---|---|---|---|---|---|
| 379 | | | lyxose | 1114-34-7 | 0.0196 | 0.0042 | 0.0019 | 0.0016 | 1.4864 | -0.3068 | -0.5722 | -0.6074 |
| 381 | | | xylose1 | 6763-34-4 | 0.0186 | 0.0188 | 0.0363 | 0.0302 | -0.8410 | -0.8197 | 1.1781 | 0.4826 |
| 385 | | | xylose2 | 6763-34-4 | 0.0984 | 0.0700 | 0.0773 | 0.0831 | 1.3427 | -1.0140 | -0.4041 | 0.0754 |
| 394 | | Amino sugar and nucleotide sugar | ribose | 24259-59-4 | 0.5027 | 0.4724 | 0.5679 | 0.5119 | -0.2774 | -1.0362 | 1.3591 | -0.0456 |
| 411 | | | ribonate, gamma-lactone | 17812-24-7 | 0.0375 | 0.0378 | 0.0417 | 0.0480 | -0.7756 | -0.6997 | 0.0971 | 1.3782 |
| 420 | | | xylitol | 87-99-0 | 0.0785 | 0.1055 | 0.0909 | 0.0726 | -0.5758 | 1.2783 | 0.2763 | -0.9788 |
| 432 | | | D-arabitol | 488-82-4 | 0.0171 | 0.0439 | 0.0174 | 0.0324 | -0.8180 | 1.2523 | -0.7952 | 0.3609 |
| 477 | | | D-glucosamine 1-phosphate | 2152-75-2 | 0.0015 | 0.0007 | 0.0067 | 0.0098 | -0.7331 | -0.9169 | 0.4640 | 1.1860 |
| 494 | | | quinate | 77-95-2 | 2.7902 | 1.2482 | 1.0448 | 1.2822 | 1.4873 | -0.4257 | -0.6781 | -0.3835 |
| 125 | | | 2-hydroxyvalerate | 617-31-2 | 0.0009 | 0.0005 | 0.0009 | 0.0006 | 0.9422 | -1.1304 | 0.7331 | -0.5449 |
| 206 | | | pelargonate | 112-05-0 | 0.0016 | 0.0024 | 0.0018 | 0.0026 | -1.0994 | 0.6565 | -0.5729 | 1.0158 |
| 354 | | | 3-hydorxy-3-methylglutarate | 503-49-1 | 0.0009 | 0.0017 | 0.0022 | 0.0033 | -1.1047 | -0.3240 | 0.1426 | 1.2862 |
| 449 | | | diglycerol | 627-82-7 | 0.0098 | 0.0096 | 0.0073 | 0.0055 | 0.8542 | 0.7617 | -0.3693 | -1.2465 |
| 551 | | | 1-hexadecanol | 36653-82-4 | 0.0005 | 0.0009 | 0.0008 | 0.0003 | -0.5501 | 1.1038 | 0.5374 | -1.0910 |
| 580 | | | palmitoleate | 373-49-9 | 0.0014 | 0.0034 | 0.0056 | 0.0052 | -1.3150 | -0.2306 | 0.8807 | 0.6649 |
| 587 | | | palmitate | 1957-10-3 | 0.5897 | 1.0433 | 0.7168 | 1.1592 | -1.0729 | 0.6197 | -0.5987 | 1.0519 |
| 603 | | Free fatty acid | linoleate methyl ester | 112-63-0 | 0.3536 | 0.2243 | 0.3974 | 0.3374 | 0.3450 | -1.4087 | 0.9386 | 0.1251 |
| 617 | | | heptadecanoate | 506-12-7 | 0.0013 | 0.0019 | 0.0036 | 0.0023 | -1.0092 | -0.3853 | 1.3533 | 0.0412 |
| 624 | | | phytol | 150-86-7 | 0.0449 | 0.0271 | 0.0000 | 0.0000 | 1.2209 | 0.4146 | -0.8178 | -0.8178 |
| 635 | | | linoleate | 60-33-3 | 0.0113 | 0.0103 | 0.0274 | 0.0216 | -0.7728 | -0.8852 | 1.1788 | 0.4792 |
| 637 | | | elaidate | 112-79-8 | 0.0035 | 0.0015 | 0.0084 | 0.0062 | -0.4706 | -1.1230 | 1.1488 | 0.4448 |
| 638 | | | linolenate | 463-40-1 | 0.1754 | 0.1395 | 0.3153 | 0.2266 | -0.5087 | -0.9790 | 1.3252 | 0.1625 |
| 644 | | | stearate | 1957-11-4 | 0.1209 | 0.1886 | 0.2020 | 0.1713 | -1.4029 | 0.5038 | 0.8820 | 0.0171 |
| 702 | Lipids | | arachidate | 506-30-9 | 0.0082 | 0.0153 | 0.0136 | 0.0177 | -1.3651 | 0.3990 | -0.0222 | 0.9883 |
| 524 | | | methyl palmitoleate | 1120-25-8 | 0.0103 | 0.0097 | 0.0148 | 0.0145 | -0.7433 | -0.9785 | 0.9224 | 0.7994 |
| 634 | | | beta-mannosylglycerate | 164324-35-0 | 0.0050 | 0.0060 | 0.0055 | 0.0092 | -0.7346 | -0.2260 | -0.5066 | 1.4672 |
| 725 | | | 2-monopalmitin | 23470-00-0 | 0.0199 | 0.0228 | 0.0218 | 0.0324 | -0.7809 | -0.2562 | -0.4266 | 1.4637 |
| 728 | | | arbutin | 497-76-7 | 0.0040 | 0.0107 | 0.0069 | 0.0112 | -1.2439 | 0.7412 | -0.3719 | 0.8746 |
| 733 | | Glycerolipids | 1-monopalmitin | 542-44-9 | 0.0110 | 0.0151 | 0.0138 | 0.0104 | -0.6849 | 1.1290 | 0.5388 | -0.9829 |
| 762 | | | 2-monoolein | 3443-84-3 | 0.0737 | 0.0298 | 0.0630 | 0.0642 | 0.8345 | -1.4527 | 0.2762 | 0.3421 |
| 773 | | | monoolein | 111-03-5 | 0.0100 | 0.0045 | 0.0071 | 0.0065 | 1.3045 | -1.1155 | 0.0397 | -0.2287 |
| 781 | | | monostearin | 123-94-4 | 0.0047 | 0.0144 | 0.0108 | 0.0072 | -1.0756 | 1.2171 | 0.3485 | -0.4901 |
| 700 | | Sphingolipid | sphingosine | 123-78-4 | 0.0047 | 0.0029 | 0.0051 | 0.0020 | 0.6937 | -0.5093 | 0.9667 | -1.1511 |
| 736 | | | phytosphingosine | 554-62-1 | 0.0045 | 0.0058 | 0.0094 | 0.0061 | -0.9398 | -0.2940 | 1.4135 | -0.1798 |
| 576 | | | Hymecromone | 90-33-5 | 0.0398 | 0.0000 | 0.0003 | 0.0000 | 1.5000 | -0.5048 | -0.4903 | -0.5048 |
| 823 | | Sterols | Stigmasterol | 83-48-7 | 0.0159 | 0.0047 | 0.0100 | 0.0086 | 1.3157 | -1.0963 | 0.0370 | -0.2564 |
| 824 | | | alpha-ecdysone | 3604-87-3 | 0.0016 | 0.0000 | 0.0000 | 0.0000 | 1.5000 | -0.5000 | -0.5000 | -0.5000 |
| 159 | | Oxidative phosphoryl | phosphate | 7664-38-2 | 0.3605 | 0.2101 | 0.2473 | 0.5695 | 0.0847 | -0.8460 | -0.6162 | 1.3775 |
| 435 | | Nicotinate and nicotin | flavin adenine degrad product(F | 146-14-5 | 0.4490 | 0.3216 | 0.3530 | 0.3058 | 1.4281 | -0.5570 | -0.0676 | -0.8035 |
| 321 | Cofactors, Prost | Ascorbate metabolism | threonate | 7306-96-9 | 0.0763 | 0.1188 | 0.0705 | 0.0674 | -0.2914 | 1.4824 | -0.5312 | -0.6598 |
| 539 | | | ascorbate | 50-81-7 | 0.0020 | 0.0007 | 0.0026 | 0.0016 | 0.3860 | -1.2790 | 1.0854 | -0.1924 |
| 583 | | | mucate | 526-99-8 | 0.0130 | 0.0167 | 0.0204 | 0.0181 | -1.2963 | -0.1262 | 1.0933 | 0.3292 |
| 153 | | Purine metabolism | oxamate | 471-47-6 | 0.0000 | 0.0039 | 0.0000 | 0.0072 | -0.7961 | 0.3140 | -0.7961 | 1.2782 |
| 194 | | | uracil | 66-22-8 | 0.0012 | 0.0053 | 0.0054 | 0.0067 | -1.4472 | 0.2680 | 0.3263 | 0.8529 |
| 245 | Nucleotides | Pyrimidine metabolism | beta-alanine | 107-95-9 | 0.0051 | 0.0226 | 0.0316 | 0.0172 | -1.2659 | 0.3107 | 1.1275 | -0.1723 |
| 705 | | | cytidine-monophosphate(CMP) | 63-37-6 | 0.0242 | 0.0211 | 0.0147 | 0.0176 | 1.1572 | 0.4123 | -1.1447 | -0.4249 |
| 706 | | | uridine | 58-96-8 | 0.0017 | 0.0002 | 0.0030 | 0.0049 | -0.3660 | -1.1320 | 0.2681 | 1.2299 |

| | | | | | | | | | | | | |
|---|---|---|---|---|---|---|---|---|---|---|---|---|
| 633 | Hormone metab | Auxin metabolism | 5-hydroxyindoleacetate | 54-16-0 | 0.0003 | 0.0009 | 0.0006 | 0.0004 | -0.9884 | 1.2544 | 0.3221 | -0.5881 |
| 42 | Secondary meta | Alkaloids | 2-Pyridinol | 142-08-5 | 0.7968 | 0.7053 | 0.7479 | 0.7482 | 1.2634 | -1.1835 | -0.0438 | -0.0360 |
| 97 | | | 3-Pyridinol | 109-00-2 | 0.0356 | 0.0204 | 0.0199 | 0.0257 | 1.4014 | -0.6874 | -0.7498 | 0.0358 |
| 101 | | | 2-piperidone | 675-20-7 | 0.0071 | 0.0065 | 0.0011 | 0.0049 | 0.8213 | 0.5865 | -1.4076 | -0.0003 |
| 163 | | | 2,3-dihydroxypyridine | 16867-04-2 | 0.0048 | 0.0023 | 0.0036 | 0.0044 | 0.9296 | -1.3555 | -0.1092 | 0.5352 |
| 185 | | | pyrrole-2-carboxylate | 634-97-9 | 0.0013 | 0.0022 | 0.0031 | 0.0034 | -1.2397 | -0.3609 | 0.6390 | 0.9616 |
| 189 | | | atropine | 51-55-8 | 0.0106 | 0.0113 | 0.0147 | 0.0149 | -1.0101 | -0.7073 | 0.8106 | 0.9068 |
| 203 | | Benzenoids | resorcinol | 108-46-3 | 0.0084 | 0.0058 | 0.0054 | 0.0072 | 1.2393 | -0.6455 | -0.9569 | 0.3631 |
| 221 | | | 3-methylcatechol | 488-17-5 | 0.0003 | 0.0166 | 0.0043 | 0.0144 | -1.0939 | 0.9814 | -0.5895 | 0.7020 |
| 232 | | | salicylate | 69-72-7 | 0.0024 | 0.0018 | 0.0044 | 0.0064 | -0.6568 | -0.9268 | 0.3144 | 1.2692 |
| 238 | | | 1,2,4-benzenetriol | 533-73-3 | 0.0026 | 0.0019 | 0.0029 | 0.0023 | 0.4692 | -1.2493 | 1.0663 | -0.2863 |
| 299 | | | phenolic phosphate | 701-64-4 | 0.0387 | 0.0417 | 0.0385 | 0.0611 | -0.5817 | -0.3051 | -0.5996 | 1.4863 |
| 356 | | | Tosylate | 104-15-4 | 0.0000 | 0.1237 | 0.2196 | 0.2379 | -1.3324 | -0.1981 | 0.6811 | 0.8495 |
| 361 | | | 4-hydroxybenzoate | 99-96-7 | 0.0069 | 0.0058 | 0.0121 | 0.0062 | -0.2828 | -0.6719 | 1.4806 | -0.5258 |
| 375 | | | gentisate | 490-79-9 | 0.0004 | 0.0021 | 0.0024 | 0.0023 | -1.4857 | 0.3215 | 0.6590 | 0.5052 |
| 377 | | | trans-3,5-dimethoxy-4-hydroxyc | 4206-58-0 | 0.0005 | 0.0085 | 0.0154 | 0.0275 | -1.0920 | -0.3903 | 0.2137 | 1.2686 |
| 459 | | | caffeate | 331-39-5 | 0.0830 | 0.0521 | 0.0866 | 0.0369 | 0.7602 | -0.5207 | 0.9103 | -1.1499 |
| 608 | | | salicin | 138-52-3 | 0.0001 | 0.0071 | 0.0031 | 0.0056 | -1.2620 | 1.0145 | -0.2931 | 0.5406 |
| 612 | | | piceatannol | 10083-24-6 | 0.0005 | 0.0000 | 0.0000 | 0.0024 | -0.2221 | -0.6253 | -0.6253 | 1.4727 |
| 618 | | | flavanone | 487-26-3 | 0.0024 | 0.0024 | 0.0057 | 0.0095 | -0.7651 | -0.7728 | 0.2056 | 1.3323 |
| 664 | | Polyketides | Butanal | 123-72-8 | 0.0016 | 0.0008 | 0.0007 | 0.0011 | 1.3651 | -0.6085 | -0.8681 | 0.1115 |
| 719 | | | DL-anabasine | 13078-04-1 | 0.0390 | 0.0328 | 0.0384 | 0.0295 | 0.8936 | -0.4720 | 0.7612 | -1.1829 |
| 724 | | Terpenoids | thymol | 89-83-8 | 0.0015 | 0.0019 | 0.0017 | 0.0022 | -1.0233 | 0.2271 | -0.4919 | 1.2881 |
